# Estimating survival of an anadromous salmonid using Bayesian hierarchical models applied to acoustic telemetry, biological, and environmental data

**DOI:** 10.1101/2024.12.04.626433

**Authors:** Inesh Munaweera, Les N. Harris, Jean-Sébastien Moore, Ross F. Tallman, Matthew J.H. Gilbert, Aaron T. Fisk, Brent G. T. Else, Mohamed M. M. Ahmed, Darren M. Gillis, Saman Muthukumarana

**Author notes:** Co-first authors (Inesh Munaweera, Les N. Harris). **Corresponding author:** Inesh Munaweera, Present Address: Mathematical and Statistical Sciences, University of Alberta, 11324 89 Ave NW, Edmonton, AB, T6G 2J5, Canada.

## Abstract

Hierarchical modelling is frequently used to model ecological processes because of its ability to handle complex ecological phenomena by decomposing them into naturally explainable sub-models. Hierarchical Bayesian approaches have gained widespread use in health, social, and environmental sciences, including in the estimation of demographic parameters such as survival. In this study, we combine Bayesian hierarchical models with acoustic telemetry data to estimate survival probabilities for high-latitude populations of an anadromous salmonid, the Arctic char *(Salvelinus alpinus)*, while incorporating environmental and biological covariates to assess their impact on survival. The model we present here can also account for temporally varying detection probabilities due to changes to the acoustic receiver array design and seasonal variation in the detection probabilities related to environmental conditions (e.g., ice vs. no ice). As previously documented in this species, survival was high (*>* 0.87) and we found that the covariates pertaining to sea ice coverage and Fulton’s condition factor impacted the survival probabilities. Contrary to our expectations, high-condition fish had lower survival rates. Survival was also considerably lower during the summer (open-water) compared to winter (ice-covered) seasons. While the biological explanations and implications of these findings require further exploration, they nonetheless demonstrate the utility of this approach. Specifically, we present a hierarchical Bayesian model that can consider environmental and biological covariates while accounting for varying detection probabilities, a major concern of acoustic telemetry studies. The model can be easily adapted for other taxa with similar life histories where mark recapture data are available and can be extended to include additional environmental (e.g., salinity) and biological parameters (e.g., sex).

## 1 Introduction

Most ecological processes are complicated and hierarchical in nature with multi-leveled spatiotemporal hierarchies (Royle and Dorazio 2009; Wikle 2003; Lele and Dennis 2009). Hierarchical modeling is frequently used to model such processes given its ability to model complex ecological phenomena by decomposing them into naturally explainable sub-models to account for different sources of variance and various sources of data which enables researchers to explore and understand the underlying mechanisms driving ecological systems (Richardson and Best 2003; Bolker 2008; Royle and Dorazio 2008). Hierarchical models can essentially be defined as the family of models that connect multiple parameters according to the structure of the problem (Gelman et al. 2004). Recently, the Bayesian approach to hierarchical models has been gaining popularity due to its extreme flexibility, better precision, and ability to incorporate prior knowledge about parameters compared to more traditional non-hierarchical models (Gelman et al. 2004; Calvert et al. 2009; Kéry and Schaub 2011). Bayesian hierarchical models extend back to the 1970s (Lindley and Smith 1972) and 1980s (Good 1980; Goel 1983) where initial applications can be found in epidemiology (Besag et al. 1991) and economics (West and Harrison 1989; Rossi et al. 1996). They are now widely applied to various types of data, including those in the financial (Chib and Greenberg 1995; Ying et al. 2005; Kantorová et al. 2020), health science (Prevost et al. 2000; Grieve et al. 2010; Kantorová et al. 2020) and environmental fields (Fuentes and Raftery 2005; Sahu et al. 2009; Apputhurai and Stephenson 2013). Bayesian hierarchical models have also been increasingly used in fisheries ecology and management for many applications such as parameter estimation for mark-recapture studies (Harley and Myers 2001; Calvert et al. 2009; Rivot and Prevost 2002), and fisheries stock assessment (Aanes et al. 2007; Aeberhard et al. 2018). More recently, combining hierarchical Bayesian models with acoustic telemetry data has facilitated the estimation of fish movement paths (Patterson et al. 2008, 2017; Munaweera et al. 2021) and other parameters of interest, such as survival probabilities and recapture rates, with better precision than traditional capture-mark-recapture models (Dudgeon et al. 2015; Alós et al. 2016; Munaweera et al. 2022).

In fisheries ecology, having a clear understanding of survival, and the biological and environmental covariates that impact survival, is vital for effective fishery conservation and management decision-making. For example, survival estimates for a particular species over time may provide insights into fishing and natural mortality, how these change over time and with changing environmental conditions. Such information allows managers to make decisions on how best to facilitate the long-term sustainability of the species (Brooks et al. 2000; Murray and Patterson 2006), for example, in setting harvest control rules (Zahner and Branch 2024). This is especially relevant for Arctic fishes whose habitats are expected to warm drastically (Rantanen et al. 2022) as a result of climate change potentially threatening survival and population persistence. Acoustic telemetry is increasingly being used to estimate demographic parameters, notably survival probability and population size (CazaAllard et al. 2021). The advantage of acoustic telemetry is that it allows for higher recapture rates and thus enhanced precision compared to traditional mark-recapture techniques (Pollock et al. 2004; Lees et al. 2021; Munaweera et al. 2022). Long-term acoustic telemetry studies, however, are hindered by the fact that acoustic telemetry arrays are often not static over the entire study period due to changing study objectives or because equipment is not recovered. Furthermore, for multi-year and multi-seasonal telemetry studies, detection probabilities vary over time, primarily due to varying environmental conditions that affect the transmission of the acoustic signals (e.g., when there is ice vs. no ice, (Munaweera et al. 2021)). This temporal variability creates problems for data analyses and interpretation of results when using traditional Cormack–Jolly–Seber (CJS) models (Cormack 1964; Jolly 1965; Seber 1965) unless varying detection probabilities are properly incorporated into the model. Anadromous (i.e., sea-run) salmonids are central to culture, food security, subsistence and health and are essential components of biodiversity and ecosystem functioning (Schtickzelle and Quinn 2007; Willson et al. 2004). They also support countless commercial and recreational fisheries significantly bolstering local economies where they occur (Galappaththi et al. 2022). Their life history also makes them vulnerable to harvest and exposes them to various anthropogenic stressors across multiple habitats (oceans, rivers and lakes) (Reist et al. 2006). Despite the importance of anadromous fishes, there is still a paucity of information on many demographic parameters in harvested stocks, and this is especially true for populations in remote Arctic locations. For example, information on survival is often lacking, and little is typically known regarding the environmental and biological variables that influence survival, and even less is known on how interannual variation in these variables affect annual survival. Evidence shows that the survival of salmonids can be influenced by water temperature (Van Wert et al. 2023; Gilbert et al. 2020). In some instances, there may also be sex-specific differences in mortality for a variety of reasons, including differences in energy exhaustion, cardiac performance, physiological stress, and immune factors between sexes (Hinch et al. 2021). Furthermore, larger individuals might have advantages in terms of predation avoidance, but they may also be preferentially selected by predators or require more resources. (Scheffer 1997; Bennett et al. 2020). This has implications for fully understanding population viability and thus, population persistence and long-term sustainability in anadromous fish stocks.

Recently, we used Bayesian multi-state capture-recapture models combined with acoustic telemetry data to estimate survival probabilities over multiple years (2014–2018) in marine/estuarine and freshwater environments for Arctic char *(Salvelinus alpinus)* in the Cambridge Bay region of Nunavut, home to Canada’s largest commercial fishery for this species (Munaweera et al. 2022). We found that marine survival was high (*>* 0.87), and survival probability was higher in freshwater compared to marine and estuary environments. Previous work in the region using capture–mark–recapture methods applied to acoustic telemetry data also revealed high survival probability in this species (0.79-0.88, Caza-Allard et al. 2021). In our previous study (Munaweera et al. 2022), however, we could only include data from receivers that were active for the entire study duration, which resulted in the exclusion of a large amount of data. Those models did not account for temporally varying detection probabilities due to seasonal factors such as surface ice cover, and we did not consider potential environmental and biological drivers that influence survival.

In this study, we extended our previous work on Arctic char by developing a more complex and flexible Bayesian hierarchical model to examine survival probabilities of anadromous fishes. The model now incorporates environmental and biological covariates to assess how variation in these impacts survival. Specifically, we included sea surface temperature (SST), surface ice conditions, Fulton’s condition factor (k) and the sex of each fish and assumed variable detection probabilities within and among years. Furthermore, we now incorporate a monthly detection variability index that was calculated using reference tag data. This factor (i.e., varying detection probability) is rarely considered in the majority of acoustic telemetry studies. Overall our aim was to develop a flexible model of survival that can account for temporal variation in array design and detection probability while incorporating additional covariates.

## 2 Materials and methods

### 2.1 Study system

Our study took place in the Cambridge Bay area of Nunavut on southern Victoria Island (Figure 1). The area is subject to long winters, and the marine environments are covered in ice from November to July. The largest commercial fishery for Arctic char in Canada takes place in the Cambridge Bay region(Harris et al. 2020a) where five waterbodies are mainly harvested with a combined annual quota of 56,100 kgs (Harris et al. 2020b). An acoustic telemetry array of 99 stations was used from 2013-2021 to study the habitat use and migrations of acoustically tagged Arctic char between marine, estuary, and freshwater environments. The acoustic array design, however, varied over time because of shifting study objectives and lost equipment. More information on the acoustic telemetry study and fishery in the region can be found in Moore et al. (2016) and Harris et al. (2020a).

**Figure 1.**
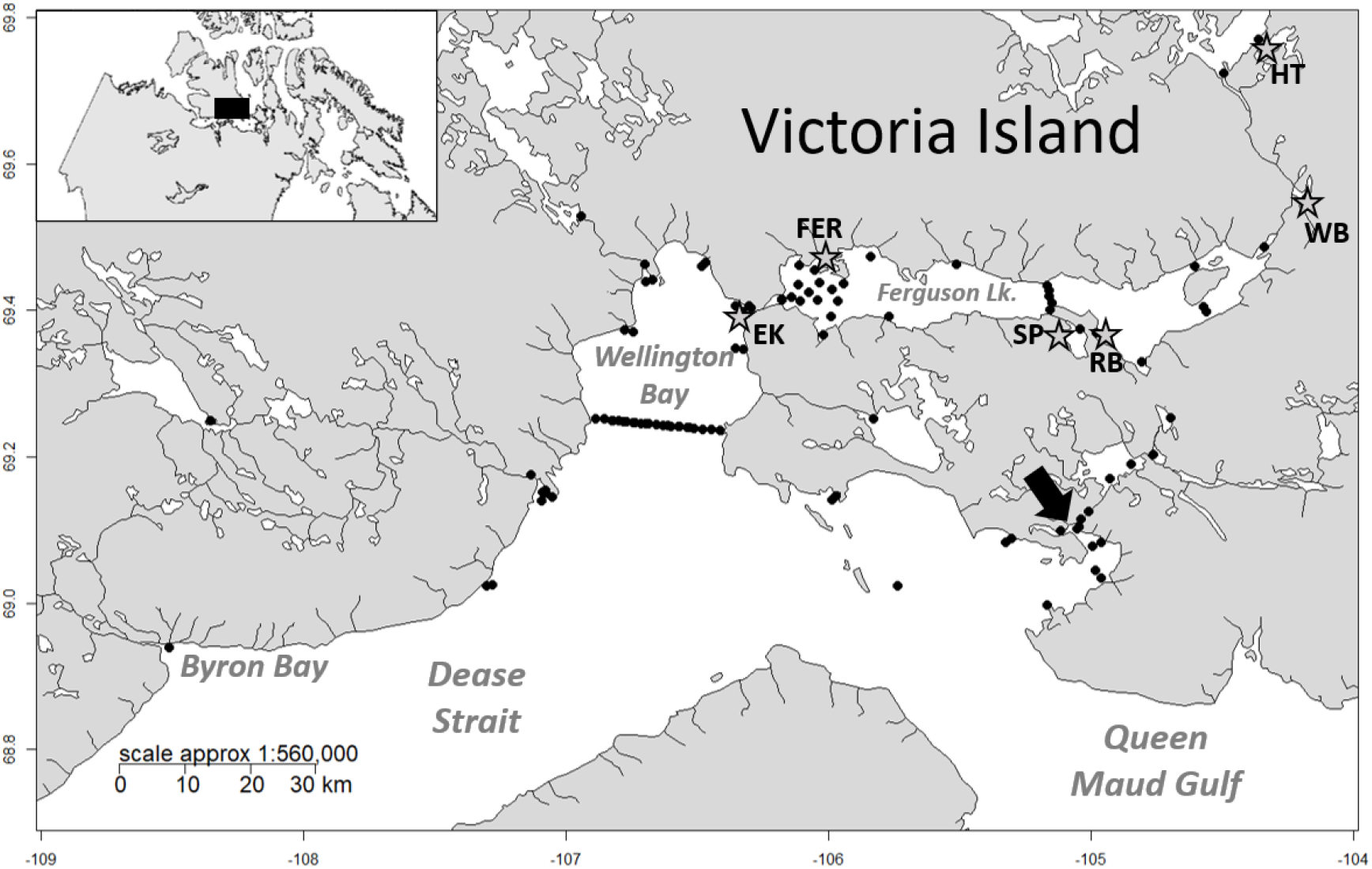
Study area on Southern Victoria Island showing all stations within our acoustic telemetry array (black dots) that were used between 2013-2020. Acoustic tagging locations are shown with stars and codes for tagging locations in this study described in Tables 1. The location of the community of Cambridge Bay is shown with a black arrow. The original map was created with R package ‘maps’ (Becker et al. 2018) using NAD83 projection, and layers for rivers and lakes were downloaded from the National Topographic Database of Canada.

### 2.2 Study design and data

This study includes data from 218 Arctic char (122 female and 96 male) that were acoustically tagged between July 2013 to August 2018 (Table 1). For each tagged Arctic char, we recorded fork length (*±*1 mm) and round weight (*±* 25 g). For each fish, Fulton’s relative condition factor (K) was calculated as *K* = [*W* × 10^5^]*/L*^3^, where W and L are weight (g) and fork length (mm), respectively. We used the genetic-sex determination protocol of Yano et al. (2013) to determine the sex of all tagged Arctic char. We estimated the daily sea surface temperature (SST) values for the marine environment in our study area from the NOAA 0.25° Daily Optimum Interpolation Sea Surface Temperature (OISST) dataset. The OISST dataset incorporates observations from different platforms (satellites, ships, buoys, and Argo floats), and uses sea ice datasets and includes a large-scale adjustment of satellite biases with respect to the in-situ data (Reynolds et al. 2007; Banzon et al. 2016; Huang et al. 2021). The dataset is interpolated using an optimum interpolation (OI) technique to fill gaps in the data and create a spatially complete map of SST over the globe. After clipping the downloaded dataset to our study area, we calculated the average monthly SST for June, July, August, and September (when Arctic char would potentially be using the marine environment) from 2013 to 2019. After September until June, Arctic char are in freshwater, but the surface temperature during this time period can safely be assumed to be less than 5^*o*^C. Weekly sea ice charts from the Canadian Ice Service (https://iceweb1.cis.ec.gc.ca/IceGraph/page1.xhtml?lang=_en) were used to determine sea surface ice conditions (SIC) for the marine environment including Coronation Gulf and Dease Strait. When the marine environment is covered in ice, we also assume that all freshwater habitats would be covered in ice. The ice cover in the marine environment is lowest in August and September, with less than 25% ice cover (Figure 3).

**Table 1.** Biological summary of Arctic char tagged in this study including tagging location (Source, EK=Ekalluk River, RB=Roberts Bay, SP=Spawning Lake, WB=Wishbone Lake, FER=Ferguson lake, HT=Heart Lake, see Figure 1), sample size (N) and weight and Fork length information.

| Year | Source | Count | Weight |  | Length |  |
| --- | --- | --- | --- | --- | --- | --- |
|  |  |  | Mean | SD | Mean | SD |
| 2013 | EK13 | 30 | 4013.3 | 1065.8 | 717.4 | 57.0 |
| 2014 | EK14 | 30 | 3156.7 | 1146.6 | 666.9 | 82.4 |
| 2015 | EK15 | 72 | 4174.3 | 1178.1 | 735.8 | 76.6 |
| 2016 | RB16 | 21 | 5319.0 | 1467.6 | 783.6 | 68.1 |
| 2016 | SP16 | 22 | 4605.7 | 1063.5 | 724.8 | 156.0 |
| 2016 | WB16 | 18 | 4244.4 | 857.1 | 734.8 | 57.7 |
| 2017 | FER17 | 6 | 4850.0 | 2911.4 | 784.0 | 81.2 |
| 2017 | HT17 | 19 | 3344.7 | 698.4 | 655.7 | 95.2 |

**Figure 2.**
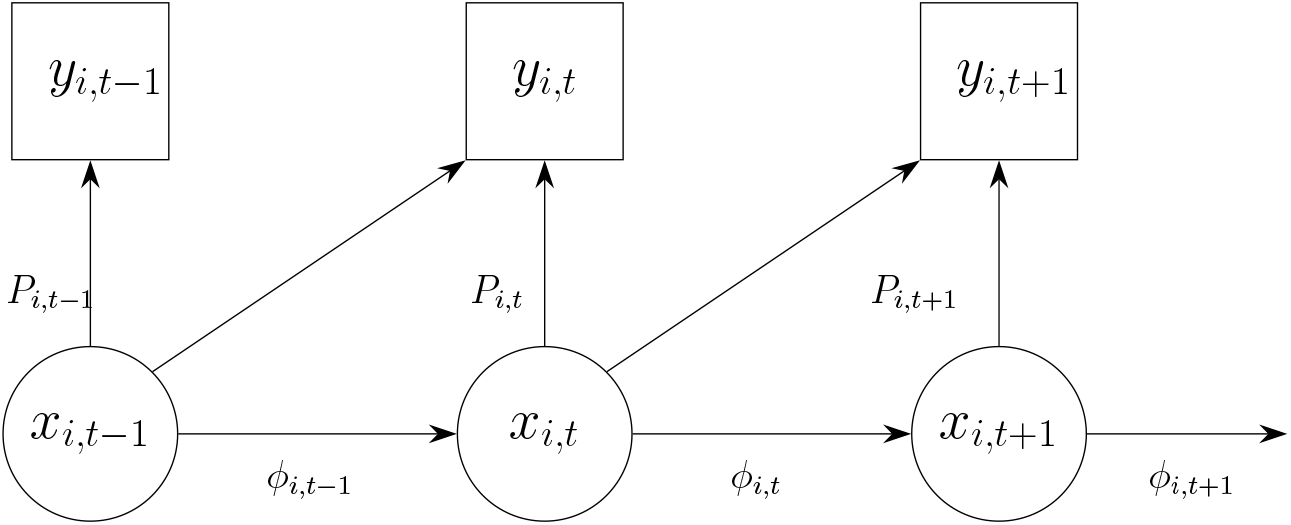
A graphical representation for the state-space formulation of the Cormack–Jolly–Seber model. Here, *x* is the latent process, *y* is the observation process, while *ϕ* and *P* are parameters that are to be estimated.

**Figure 3.**
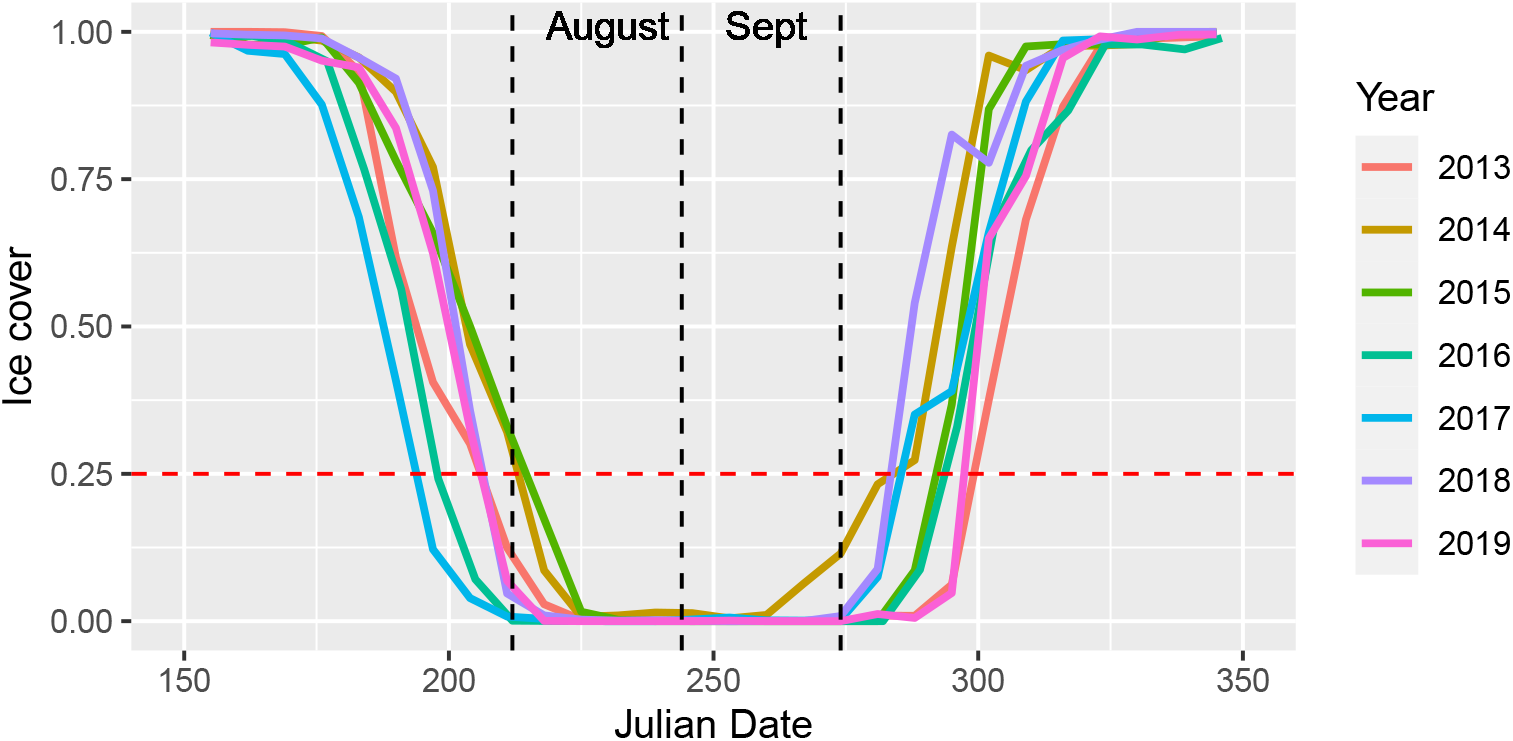
Weekly sea ice cover from 2013 to 2019.

### 2.3 Bayesian hierarchical models

Bayesian hierarchical modeling is a key framework in Bayesian statistics that allows for the modeling of obscure population dependencies using a much simpler set of layers to represent different levels of hierarchy (Gelman et al. 2013; Shaddick and Zidek 2015). The state-space model is one of the most popular hierarchical models that has three layers: 1) the observation model that describes observed data generated from a latent process (complete data likelihood), 2) the process model that represents the underlying latent process (latent data density), 3) the prior densities (model for the parameters) based on prior knowledge (Shaddick and Zidek 2015; Congdon 2010). The popular Cormack–Jolly–Seber (CJS) model (Cormack 1964; Jolly 1965; Seber 1965) can be easily formulated using state-space formulation (King 2012; Kéry and Schaub 2011). Figure 2 shows a graphical representation of the state-space formulation of the CJS model.

Consider a regular mark-recapture setting with *n* number of tagged fish observed through *T* number of recapture instances. Then, the joint density can be generally expressed below (Congdon 2010; Fouley 2013; Kéry and Schaub 2011).

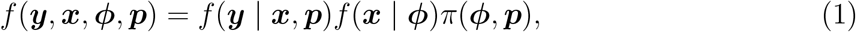

where *f* (***y*** | ***x, p***) is the observation model for the observed data (***y***). The observation model is conditioned on the latent (hidden) states of captured fish (***x***) and the detection probability (***p***). Note that ***x*** and ***y*** are *n* × *T* matrices where the elements of each matrix are defined as:

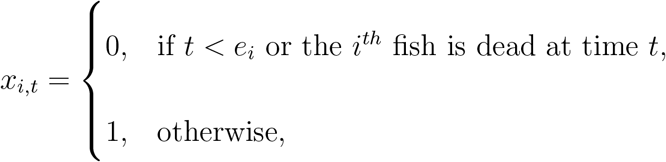

where *e*_*i*_ is the tagged time step of the *i*^*th*^ fish, and

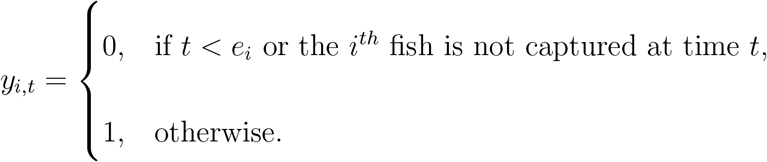

Then, the observation model can be written as

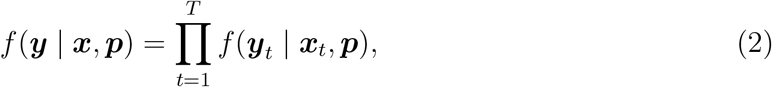

where ***y***_*t*_ = {*y*_1,*t*_, *y*_2,*t*_, …, *y*_*n,t*_} and ***x***_*t*_ = {*x*_1,*t*_, *x*_2,*t*_, …, *x*_*n,t*_} are the vector of observed status and the vector of latent states respectively at time step *t* ∈ {1, 2, …, *T*}.

The second part of the joint density in (1), *f* (***x*** | ***ϕ***) is the process model that describes the latent states (***x***) of the fish that depend on the survival probability (***ϕ***) given by

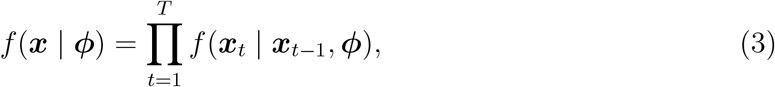

The third component of the joint density, *π*(***ϕ, p***) is the joint prior density of the parameters ***p*** and ***ϕ*** (Congdon 2010; Shaddick and Zidek 2015).

The posterior density can be written as

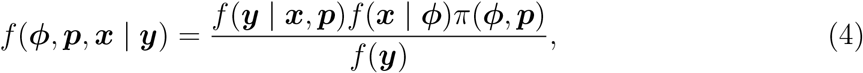

where, *f* (***y***) is called the normalizing constant. The marginal posterior distribution for the parameters ***ϕ, p*** can be obtained by

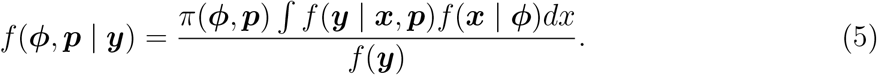

The posterior distribution in Eq. 5 is obtained indirectly by simulating samples using the Bayesian Markov chain Monte Carlo (MCMC) approach since the analytical solution for the posterior distribution is intractable (Congdon 2010; King 2012). In the MCMC approach, we use the much simpler conditional distributions *f* (***ϕ, p*** | ***x, y***) and *f* (***x*** | ***ϕ, p, y***) to fully characterize the posterior in Eq. 5. In the *k*^*th*^ iteration, MCMC will alternatively sample from the conditional distributions *f* (***x***^(*k*)^ | ***ϕ***^(*k*−1)^, ***p***^(*k*−1)^, ***y***) and *f* (***ϕ***^(*k*)^, ***p***^(*k*)^ | ***x***^(*k*)^, ***y***) (Congdon 2010; King 2012).

The process model in Eq. 2 is defined by a Bernoulli distribution given by

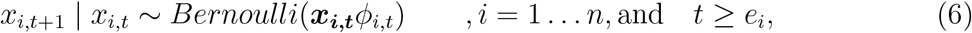

where *e*_*i*_ is the tagged time step of the *i*^*th*^ fish. Similarly, the observation model in Eq. 3 is defined as

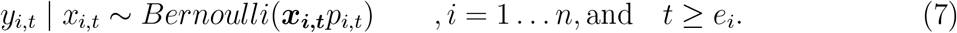

We will assume that the survival probability *ϕ*_*i,t*_ is associated with the temporal covariates: sea surface temperature (SST) and surface ice condition (SIC), and individual covariates; Fulton’s condition factor (FC) and sex. Also, we will assume an individual random effect (*ϵ*_*i*_) to account for any remaining individual variability (Kéry and Schaub 2011). Then, we will model the survival as below

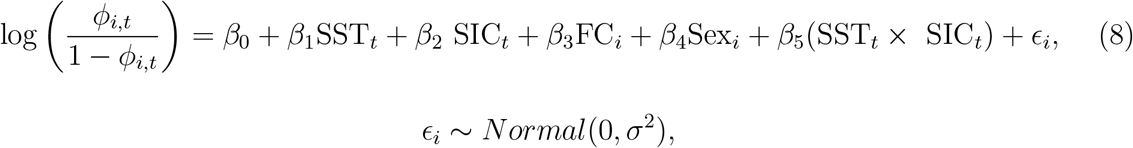

where the SST is obtained by taking the average sea surface temperature for the month as below 5C^°^ (SST=0) or above 5C^°^ (SST=1) to assess if SST impacts survival at lower and higher values, we were restricted to lower values of temperature given that the highest monthly average temp was 8C^°^. The threshold of 5C^°^ was chosen based on Arctic char preferred temperatures being between 5C^°^ and 8C^°^ (Harris et al 2020), however, this model can be extended to any range of temperatures the study organism would experience in nature. Fulton’s condition factor was recorded as below 0.9 (FC=0) or above 0.9 (FC=1) to assess if there were survival differences between fish with lower and higher FC when they were acoustically tagged. Finally, we also recorded the marine surface ice condition as a binary variable that reflects whether the surface is covered in ice (SIC=0, if the marine ice cover is *>*25%) or mostly open water (SIC=1, otherwise) during the given month to assess the potential impact of surface ice conditions on survival (Figure 3). Based on previous work in the region, low K (less than 0.7) is thought to be associated with elevated at-sea mortality (Dutil, 1986). There were limited individuals with very low K values in our dataset, so we chose a higher threshold to still allow us to test the utility of incorporating K as a covariate influencing survival. As with SST, any range of K could be explored with this model if data were available.

Since the detection probabilities exhibited monthly variation within and among years, we used a monthly detection variability index (DVI) calculated with reference tag data for each habitat separately (marine, estuary, and freshwater) to account for the within-year variability in the detection probabilities. Each reference tag was approximately 500 m away from the closest receiver, and any missing values during the study period were imputed with the monthly averages across years. For each fish, the DVI at time *t* was assigned according to the location of the most recent detection record before time *t*. Note that DVI is just an indicator of the variability in the detection probabilities within a year for each environment. The exact probability that a fish would be detected by at least one of the receivers during the given period depends on how far the fish is from the nearest receiver, how many receivers were active during the period and other environmental conditions (e.g., ice and wind). DVI can account for the monthly variability in detection probabilities due to these environmental variables that change seasonally and spatially (since we calculate DVI for each environment). A DVI above 1 indicates above-average detection probability, and below one indicates lower than average detection probability. Our acoustic array changed annually by adding new receivers, removing receivers and/or repositioning current receivers. Hence, we assume annually varying detection probabilities to account for those changes to the receiver array. Furthermore, tags were programmed to automatically turn off at 72 months. Hence, the detection probability was assigned zero for all the fish once the time from tagging was greater than 72 months. Then we modelled the detection probability as below:

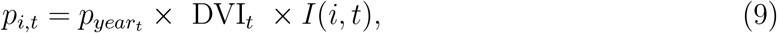

where *p*_*year*_ is the average detection probability for the year of month *t* which is unknown. Here, *I*(*i, t*) is a indicator function such that

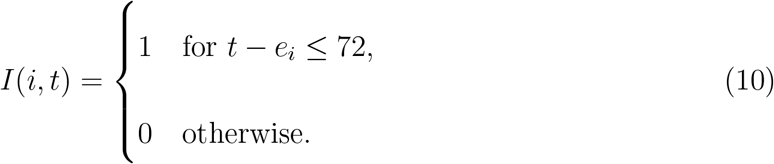

Note that the receiver array has been changed annually; hence we have to estimate the average detection probabilities for each year, given by *p*_*year*_.

#### 2.3.1 Data preparation

In our study, the fish are detected by an array of receivers, and the transmission of an acoustic signal from a fish to a receiver takes just a few milliseconds; each detection can be considered as an instantaneous sampling point. There were millions of such detection records in our data set. Hence, we pooled the detection record monthly. If the fish was detected at least once during the month, we recorded the fish as alive during that month.

### 2.4 Model estimation and evaluation

The models were fitted using the Bayesian MCMC approach with ‘JAGS’ in R using the package ‘R2jags’ (Su and Yajima 2015). The models were run on Compute Canada, Beluga cluster that uses 2×Intel Gold 6148 Skylake @ 2.4GHz processors with 40 cores in each node. For each model, three parallel chains were run with 100,000 MCMC iterations and thinning was done by selecting each 5^*th*^ iteration of the MCMC chain to minimize any possible autocorrelations in the simulated parameter values. We used the Effective Sample Size (ESS) suggested by Kass et al. (1998) and Gelman-Rubin statistics 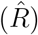 by Gelman and Rubin (1992) to assess convergence. ESS can be used to measure the amount of independent information in the MCMC chain. We can calculate 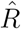 for the parameter of interest by dividing the total variance of the parameter of interest calculated with multiple MCMC chains combined by the average of the variances within each chain (Kass et al. 1998). Hence, a 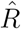 close to 1 denotes a converged MCMC chain for the parameter.

To check for any identifiability issues, we inspected the posterior distributions, trace plots and pairwise correlations between the parameters generated with MCMC chains. An identifiable model should result in unimodal posterior distributions, trace plots of multiple chains with good mixing (Siekmann et al. 2012; Simpson et al. 2020) and weak correlations between model parameters (Hines et al. 2014).

### 2.5 Model selection

To perform the model selection, we used the deviance information criterion (DIC) proposed by Spiegelhalter et al. (2002). DIC accounts for both the model fit and the complexity of the model (Gelman et al. 2013). DIC can be easily obtained using the Bayesian programming language `‘JAGS’` (Plummer 2003; Spiegelhalter et al. 2003). In this work, we used the version of DIC suggested by Gelman et al. (2004) given below:

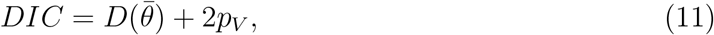

where 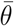 is the posterior mean and *p*_*V*_ is given by

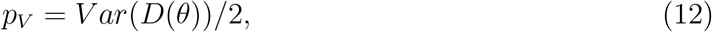

and

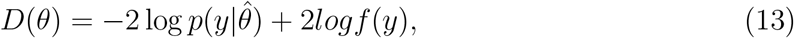

where, *f* (*y*) is the standardizing term that is a function of data (Spiegelhalter et al. 2002).

## 3 Results

The monthly detection index exhibited a regular pattern with below-average detection probabilities in summer and above-average detection probabilities in winter (Figure 4). This behaviour was similar for all three environments, but the variability was comparatively lower in the estuary environment and higher in freshwater. In the marine environment, we observed a very noticeable drop in the detection probability, with the minimum around September compared to other environments and a higher, almost constant detection probability from December to June.

**Figure 4.**
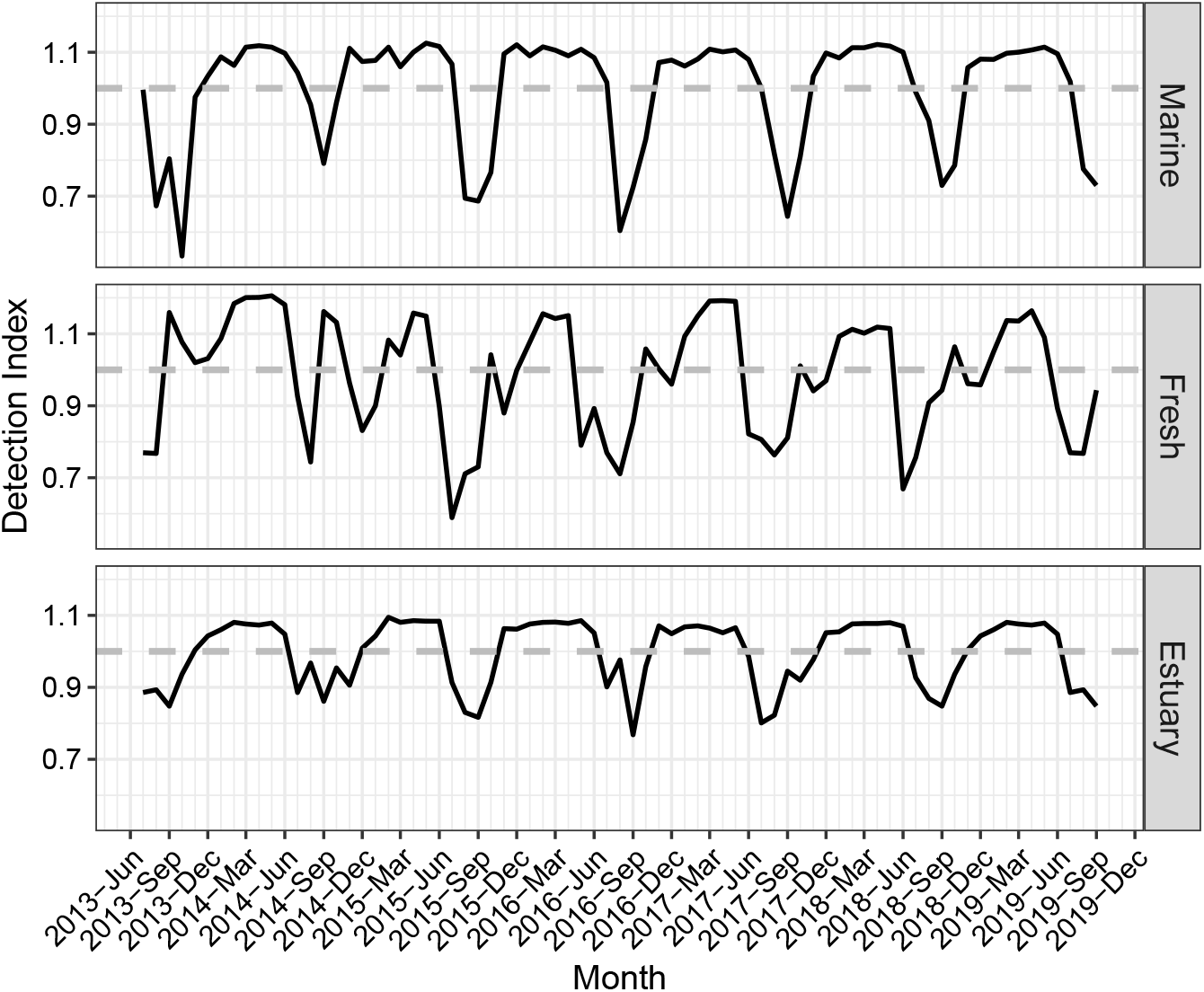
Monthly detection variability index (DVI) shown for marine, freshwater and estuarine habitats incorporated in this study (from June-2013 to December 2019). Horizontal lines on each plot represent the average DVI (note that DVI has been scaled so that average DVI = 1).

The full model using all covariates in the survival probability function in (8) did not converge, and hence, we dropped the individual random effect (*ϵ*_*i*_) in all models. The best model with the lowest DIC was the model with surface ice condition and Fulton’s condition factor (Table 2). The model did not suffer from convergence issues, confirmed by Gelman-Rubin statistics of almost one and high ESS values (≥ 1900) for all parameters. The model also did not suffer from any identifiability issues since all the posterior densities were unimodal, and the trace plots showed multiple chains mixed well for all the parameters. The 95% credible intervals for all the estimated coefficients for the best model did not contain zero, suggesting that all the coefficients in the model are relevant in predicting the survival of Arctic char (Table 3). The estimated annual detection probabilities were precise, with low standard errors.

**Table 2.** DIC values for fitted models with different covariates for the survival model. The best model is the one with the lowest DIC (marked by ‘*’). ΔDIC is the difference in the DIC scores from best model. The covariates are sea surface temperature (SST), surface ice condition (SIC), Fulton’s condition factor (FC), and sex.

| Survival covariates | DIC | ΔDIC |
| --- | --- | --- |
| SST,Sex,FC,SIC,(SST×SIC) | 8601.0 | 12.2 |
| SST,FC,SIC,(SST×SIC) | 8601.1 | 12.3 |
| SST,FC,SIC | 8604.0 | 15.2 |
| Sex,FC,SIC | 8600.6 | 11.8 |
| FC,SIC | 8588.8* | 0 |

**Table 3.** Parameter estimates for the best model with the lowest DIC. Here, *β*_0_, *β*_2_ and *β*_3_ represent the intercept, coefficients for sea surface condition, and Fulton’s condition factor, respectively. *p*_1_, *p*_2_, …, *p*_7_ represents the annual detection probabilities. The effective sample size is denoted by ESS, and 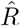 denotes the Gelman-Rubin statistics.

| | Mean | SE | Quantiles | | $\hat{R}$ | ESS |
| --- | --- | --- | --- | --- | --- | --- |
|  |  |  | 2.5% | 97.5% |  |  |
| $\beta_0$ | 5.897 | 0.366 | 5.226 | 6.661 | 1.002 | 1900 |
| $\beta_2$ | -3.555 | 0.338 | -4.263 | -2.937 | 1.002 | 2300 |
| $\beta_3$ | -0.477 | 0.239 | -0.966 | -0.028 | 1.001 | 9700 |
| $p_1$ | 0.286 | 0.038 | 0.216 | 0.363 | 1.001 | 18000 |
| $p_2$ | 0.222 | 0.021 | 0.183 | 0.264 | 1.001 | 32000 |
| $p_3$ | 0.532 | 0.017 | 0.497 | 0.566 | 1.001 | 25000 |
| $p_4$ | 0.540 | 0.013 | 0.513 | 0.565 | 1.001 | 18000 |
| $p_5$ | 0.517 | 0.012 | 0.493 | 0.541 | 1.001 | 27000 |
| $p_6$ | 0.421 | 0.015 | 0.392 | 0.449 | 1.001 | 33000 |
| $p_7$ | 0.236 | 0.019 | 0.199 | 0.275 | 1.001 | 11000 |
| Deviance | 8201.5 | 27.8 | 8148.641 | 8257.8 | 1.001 | 12000 |

The marginal probability densities for the survival probabilities of Arctic char showed a clear difference between levels of surface ice condition and Fulton’s condition factor (Figure 5). The survival probability of Arctic char was significantly lower in the open water season (when there was less ice) compared to the ice-covered season. Furthermore, we also notice slightly lower survival for Arctic char with higher Fulton’s condition factor, which was particularly pronounced in the open water season (Table 4). Finally, the average monthly survival estimates obtained by averaging the simulated survival estimates of the MCMC output showed a noticeable drop in the survival probability of Arctic char in July and August (Figure 6).

**Table 4.** Summary of marginal survival probabilities for different levels of surface ice condition and Fulton’s condition factor. The credible intervals were obtained using the 2.5%^*th*^ percentile and 97.5%^*th*^ percentile of each marginal density extracted using the MCMC outputs.

|  |  | Surface condition |  |
| --- | --- | --- | --- |
|  |  | Mostly ice covered | Mostly open water |
| Fulton’s conditon factor | <0.9 | Median = 0.997 | Median = 0.912 |
|  |  | 95% CI = (0.995,0.999) | 95% CI = (0.872,0.943) |
|  | >0.9 | Median = 0.996 | Median = 0.866 |
|  |  | 95% CI = (0.992,0.998) | 95% CI = (0.841,0.888) |

**Figure 5.**
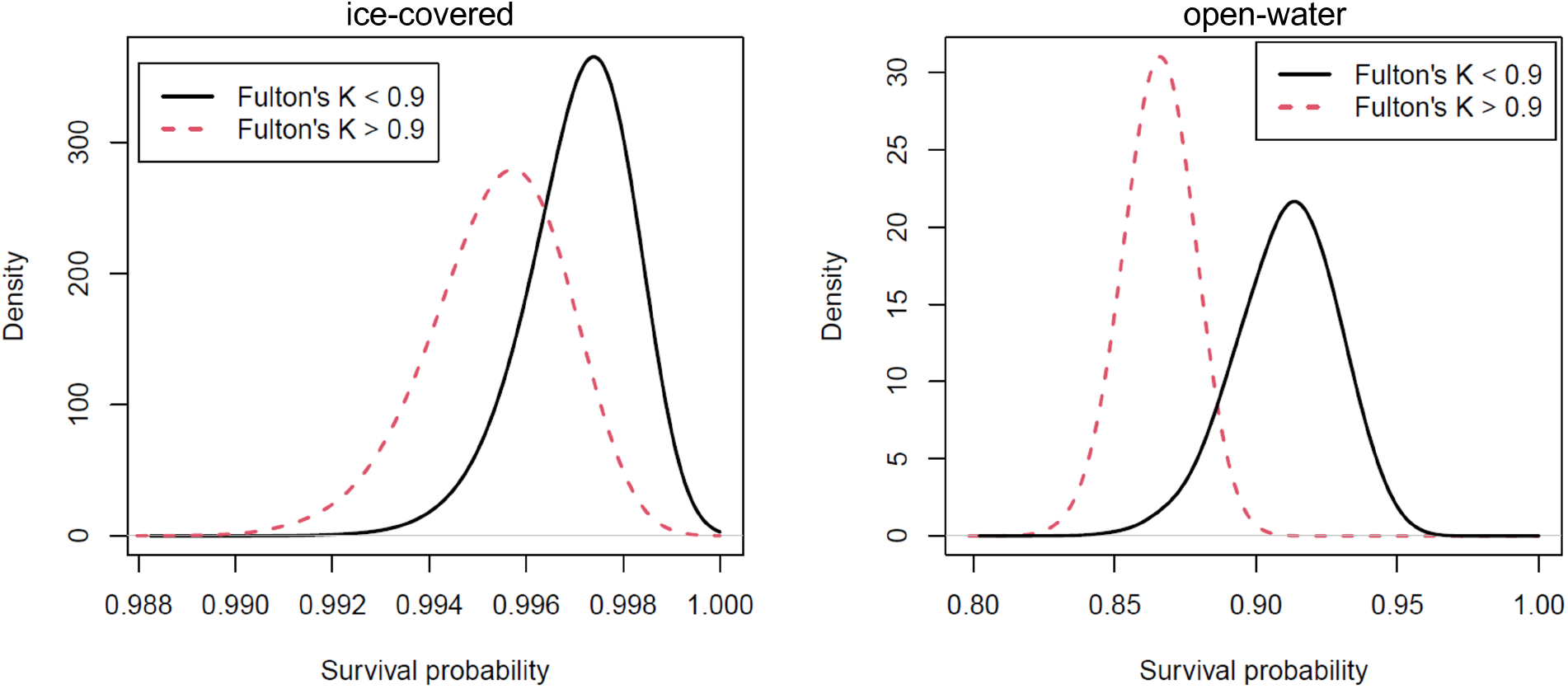
Marginal densities of survival probabilities for different levels of Fulton’s condition factor and surface ice condition (left - ice-covered, right - open water). All the densities were obtained using kernal density estimation with the Gaussian kernal with arbitrarily selected bandwidths of 0.0005 (left) and 0.005 (right).

**Figure 6.**
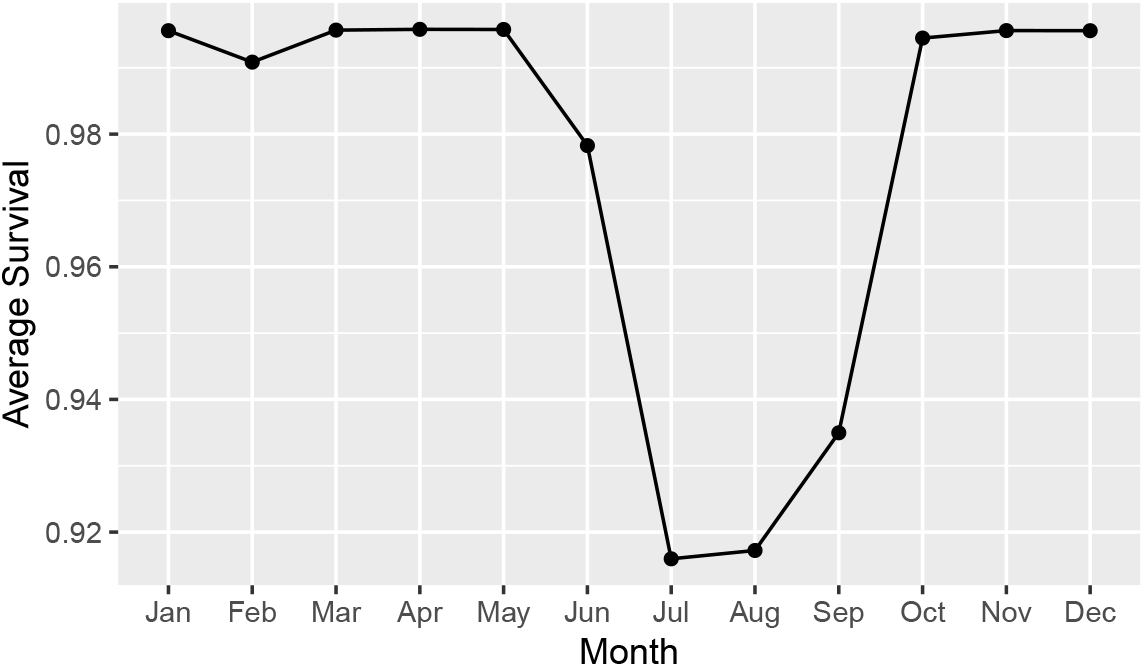
Monthly survival estimate obtained by averaging the simulated survival estimates using MCMC outputs.

## 4 Discussion

Disentangling the impacts of environmental and climatic variables on the survival of anadromous and freshwater fishes provides information that is critical for predicting the effects of climate change on population resilience (Reist et al. 2006; Crossin et al. 2017; Caza-Allard et al. 2021). Furthermore, understating the biological drivers impacting survival has implications for understanding population persistence and long-term sustainability. However, formulating models for the survival of this group of fishes is often challenging because many methods (i.e., those based on age) require fish to be sacrificed, or there is a paucity of data for estimating these demographic parameters. This issue is particularly pronounced in Arctic study systems. Analyses of acoustic telemetry data have recently provided alternative approaches for estimating survival in various taxa; however, covariates, non-static array design and varying detection probabilities have rarely been considered in such models. Here, we present a hierarchical Bayesian approach for estimating survival over seven years in Arctic char from the Cambridge Bay region of Nunavut. Our model incorporates environmental (sea surface temperature and surface ice condition) and biological (Fulton’s condition factor and sex) covariates as well as varying detection probabilities that account for the changes in the acoustic array from year to year. We found that survival of Arctic char was high across all years, and the surface ice condition and Fulton’s condition factor influence the survival probability of Arctic char in the region. The models converged well (three chains each with 100,000 iterations after 20,000 burn-in) without any identifiability issues. To our knowledge, this study was the first to model survival using Bayesian models and acoustic telemetry data with environmental and biological covariates while considering within-year and between-years varying detection probabilities.

Modeling demographic parameters using the Bayesian hierarchical framework is gaining popularity because of its flexibility, ability to incorporate prior knowledge, and the parameter estimates with the Bayesian hierarchical approach frequently yield better precision and lower bias than traditional non-hierarchical models (Clark 2005; Calvert et al. 2009). For example, Gimenez et al. (2007) used a Bayesian state-space modeling approach with CJS models to estimate survival of the European dipper (*Cinclus cinclus*). Poole (2002) utilized a Bayesian framework to estimate survival rates of fulmar petrels (*Fulmaris glacialis*) in Orkney, leveraging 13 years of mark-recapture data. These studies showcased the Bayesian approach’s capability to adeptly manage missing data, demonstrating its utility in addressing gaps within longitudinal datasets. Although the Bayesian approach incorporating acoustic telemetry data is gaining popularity (e.g., Lees et al. 2021), studies that incorporate environmental and biological covariates and address issues with seasonally and annually varying detection probability remain rare.

Caza-Allard et al. (2021) previously used the CJS models with maximum likelihood method to understand the influence of the environmental and biological covariates on the Arctic char survival in our study area. However, their study was limited to the marine environment, and they did not consider the within-year variation in survival and detection probabilities. Their most parsimonious model (assessed using QAICc) included the effect of sea ice melt date, however, the value of this parameter did not differ from zero, suggesting further study is needed to resolve its potential importance in influencing survival. The hierarchical Bayesian framework based on the CJS we presented here is an effective method for estimating survival in a study system that includes multiple populations, multiple years of data, annual differences in array design, tag characteristics and highly variable detection probabilities. Additionally, this model can be extended to other acoustic telemetry data sets where environmental and biological covariates are available and where detection probabilities vary spatially and temporally within the acoustic array.

In this study, we incorporated covariates in survival estimation and found that the surface ice condition and Fulton’s condition factor (K) impacted survival probability of Arctic char. Specifically, survival was lower in the open-water season and in Arctic char with higher K. Although we only know K at the time of tagging and cannot follow K across years, this model would still be valuable for short-term studies where K is known and where there are large differences in K among individuals. In general, the average K is typical for healthy anadromous salmonids (mean=1.02 and SD = 0.27, e.g., Gilbert et al. (2016)), and we assume that K would only increase through the open water feeding season when fishing pressure would be the highest. Thus, the high condition of fish in this study likely did not increase mortality for biological reasons (i.e., fish health). Rather, fish with a higher K (those that are plumper) may have been more susceptible to the commercial and subsistence harvest that occurs in our study area. Indeed, acoustic telemetry and other mark-recapture initiatives have revealed that acoustically tagged Arctic char are occasionally harvested in these fisheries (Harris et al. 2022). Large and better-condition fish may also be more susceptible to predation as seals, loons, gulls, and narwhal *(Monodon monoceros)* are all known to feed on char in the region (Gilbert et al. in review). Finally, larger char may be more prone to stranding-related mortality during their upriver migration when water levels are commonly low (Gilbert et al. 2016). We found that neither sex nor SST influenced survival. This might be mainly due to the low variability of sea surface temperature having an effect on the survival of Arctic char. The maximum average monthly temperature reported in the region for the study duration is only 8 C^*o*^, which might not be detrimental to Arctic char survival (Harris et al. 2020a; Gilbert et al. 2020). Furthermore, this may also reflect a limitation in the SST data; the low spatial resolution product is likely incapable of capturing SST variability in shallow, near-shore environments like estuaries. We found no difference in survival between sexes and others have also shown that mortality between sexes is similar for Arctic char in the area (Zhu et al. 2017, 2021). This finding is in contrast to some other anadromous salmonid systems, which can exhibit a strong female-biased mortality (Hinch et al. 2021)

In multi-year acoustic telemetry studies, the acoustic array is often not consistent over the study period for a variety of reasons (objectives change, lost receivers, etc.). Additionally, detection probabilities are not static and change throughout the study (within and among years) depending on the habitat occupied (e.g., riverine vs. lacustrine and temporally varying environmental conditions(e.g., ice vs. no ice)). Despite having an acoustic array that varied annually and detection probabilities that changed monthly and with the environmental conditions of the habitats Arctic char were using, we could incorporate varying detection probabilities and stochastic array design in our model. This approach allowed us to incorporate all data collected throughout the study period instead of ignoring the data collected by the receivers that were lost, removed or re-positioned during the study period resulting in more precise estimates. Hence, the modeling framework we developed here would be useful for any telemetry study where we know that the detection probabilities change over time and where there are temporal changes in the array design, which is a common feature of acoustic telemetry studies.

In the model estimation, we had to drop the individual variability term in the survival model due to convergence issues. We believe that this might be due to overparameterization that occurred by adding the additional error component to each fish. We may be able to eliminate this as we collect more data that are not available at the moment. Additional information that might further improve the model is fishing effort information, predator densities and food availability among seasons and habitats and knowledge of migratory conditions in the region (Gilbert et al. 2020). A detailed examination of how additional environmental (e.g., salinity) and biological covariates (e.g., age) influence Arctic char survival remains warranted and necessary for further understanding of how anadromous fishes, in general, persist and survive as they encounter multiple habitats that vary substantially.

## 5 Conclusions

In this study, we used Bayesian hierarchical models with acoustic telemetry data to estimate survival probabilities for anadromous Arctic char incorporating environmental and individual biological covariates to assess the impact of these factors on survival in this species. The model also accounted for temporally varying detection probabilities, which is often not considered in acoustic telemetry studies. The results of this work should prove valuable for furthering our collective understanding of the environmental and biological drivers of survival in anadromous Arctic char and salmonids in general. Additionally, the proposed approach is very flexible and can be adapted to study other taxa. For instance, the proposed Bayesian hierarchical modeling framework can be easily modified to study the survival of animals monitored with a grid of automated observer stations that detect the animal’s presence with detection probability affected by temporal and spatial factors. The proposed model can also account for any spatial and temporal changes in the receiver grid design. Two examples are camera traps with motion sensors and radio tags to study wildlife survival (Caravaggi et al. 2020; Griffin et al. 2019). Finally, the Bayesian hierarchical modeling framework is an extremely flexible and powerful tool that can be used to effectively model complex error structures of ecological data collected using modern animal tracking technologies. With ongoing advancements in computing power and more efficient MCMC samplers, we are certain that the Bayesian hierarchical modeling framework will be increasingly used as the natural candidate for modeling such data, including acoustic telemetry data used to estimate demographic parameters.

## 6 CRediT authorship contribution statement

**Inesh Munaweera:** Conceptualization, Data curation, Formal analysis, Investigation, Methodology, Visualization, Writing – original draft, Writing – review & editing. **Les Harris:** Conceptualization, Funding acquisition, Data curation, Investigation, Methodology, Project administration, Resources, Supervision, Writing – original draft, Writing – review & editing. **JS Moore:** Resources, Funding acquisition, Data curation, Investigation, Methodology, Project administration, Writing – review & editing. **Ross Tallman:** Resources, Writing – review & editing. **Matt Gilbert:** Resources, Investigation, Writing – original draft, Writing – review & editing. **Aron Fisk:** Resources, Funding acquisition, Project administration, Resources, Writing – review & editing. **Brent Else:** Resources, Writing – review & editing. **Mohamed Ahmed:** Resources, Investigation, Writing – review & editing. **Darren Gillis:** Conceptualization, Funding acquisition, Methodology, Project administration, Resources, Supervision, Writing – review & editing. **Saman Muthukumarana:** Conceptualization, Funding acquisition, Methodology, Project administration, Resources, Supervision, Writing – review & editing.

## 8 Data Availability

All data and code used in this article can be made available upon request.

## Acknowledgment

We thank the countless field personnel who made this work possible. Specifically, we thank M. Omilgoetok, R. Ekpakohak, D. Kanayok and K. Kanayok. Field support was also provided by B. Malley and M. Falardeau. We owe extended thanks to the Ekaluktutiak Hunters and Trappers Organization for support of this project and for logistic support during the field component of this study. Funding and/or logistical support was provided by the Ocean Tracking Network, Fisheries and Oceans Canada (Nunavut Implementation Funds), the Nunavut Wildlife Management Board, Polar Knowledge Canada, the Marine Environmental Observation Prediction and Response Network, Arctic Net, Dal Aviation, the Polar Continental Shelf Project and the Arctic Research Foundation crew of the RV ‘Martin Bergmann’. Dr. Muthukumarana and Dr. Gillis have been partially supported by discovery grants from the Natural Sciences and Engineering Research Council of Canada. We thank the associate editor and XXX anonymous reviewers for constructive feedback that improved earlier versions of the manuscript.

## Appendix

**Figure A1.**
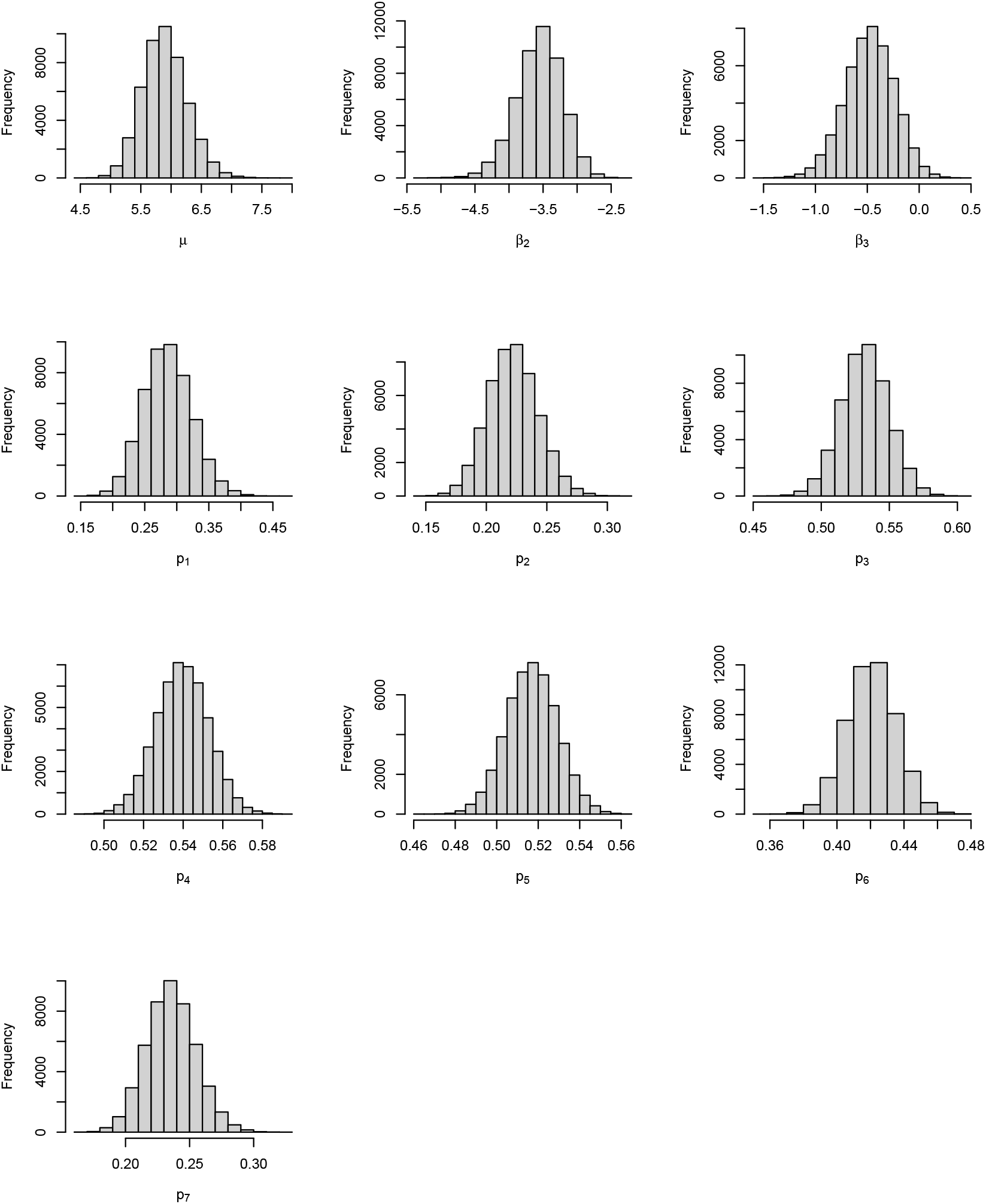
Posterior plots of parameter estimates for the best model. Here, *β*_0_, *β*_2_ and *β*_3_ represent the intercept, coefficients for sea surface condition, and Fulton’s condition factor respectively. *p*_1_, *p*_2_, …, *p*_7_ represents the annual detection probabilities.

**Figure A2.**
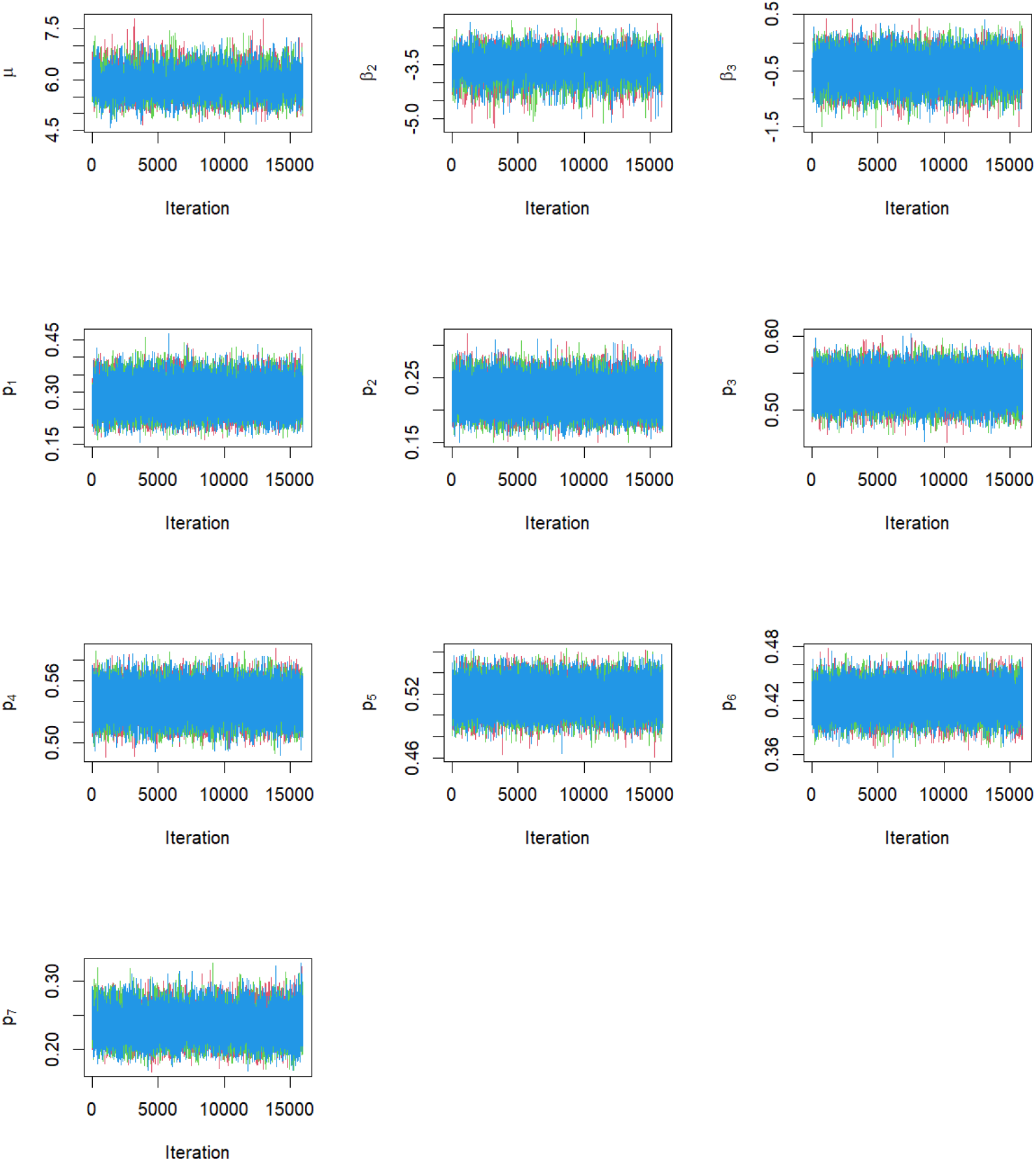
Trace plots of parameter estimates for the best model. Here, *β*_0_, *β*_2_ and *β*_3_ represent the intercept, coefficients for sea surface condition, and Fulton’s condition factor respectively. *p*_1_, *p*_2_, …, *p*_7_ represents the annual detection probabilities.

